# A Positive Feedback Loop Ensures Propagation of ROS Production and JNK Signaling Throughout *Drosophila* Tissue Regeneration

**DOI:** 10.1101/152140

**Authors:** Sumbul Jawed Khan, Syeda Nayab Fatima Abidi, Andrea Skinner, Yuan Tian, Rachel K. Smith-Bolton

## Abstract

Regenerating tissue must initiate the signaling that drives regenerative growth, and sustain that signaling long enough for regeneration to complete. How these key signals are sustained is unclear. To gain a comprehensive view of the changes in gene expression that occur during regeneration, we performed wholegenome mRNAseq of actively regenerating tissue from damaged *Drosophila* wing imaginal discs. We used genetic tools to ablate the wing primordium to induce regeneration, and carried out transcriptional profiling of the regeneration blastema by fluorescent labeling and sorting the blastema cells, thus identifying differentially expressed genes. Importantly, by using genetic mutants of several of these differentially expressed genes we have confirmed that they have roles in regeneration. Using this approach, we show that high expression of the gene *moladietz* (*mol*), which encodes the Duox-maturation factor NIP, is required during regeneration to produce reactive oxygen species (ROS), which in turn sustain JNK signaling during regeneration. We also show that JNK signaling upregulates *mol* expression, thereby activating a positive feedback signal that ensures the prolonged JNK activation required for regenerative growth. Thus, by wholegenome transcriptional profiling of regenerating tissue we have identified a positive feedback loop that regulates the extent of regenerative growth.

**Author summary:** Regenerating tissue must initiate the signaling that drives regenerative growth, and then sustain that signaling long enough for regeneration to complete. *Drosophila* imaginal discs, the epithelial structures in the larva that will form the adult animal during metamorphosis, have been an important model system for tissue repair and regeneration for over 60 years. Here we show that damage-induced JNK signaling leads to the upregulation of a gene called *moladietz*, which encodes a co-factor for an enzyme, NADPH dual oxidase (DUOX), that generates reactive oxygen species (ROS), a key tissue-damage signal. High expression of *moladietz* induces continuous production of ROS in the regenerating tissue. The sustained production of ROS then continues to activate JNK signaling throughout the course of regeneration, ensuring maximal tissue regrowth.

## Introduction

The capacity to regenerate damaged or lost organs or limbs is significantly greater in some animals than others. The use of model organisms with varying degrees of regenerative capacity, from whole-body regeneration in planaria and hydra, to limb regeneration in amphibians, organ and fin regeneration in zebrafish, and the limited tissue regeneration that occurs in mammalian models, has advanced our understanding of this process (reviewed in 1). The complementary tools available in different model organisms has enabled identification of conserved mechanisms and signaling pathways that are used in many regeneration contexts, such as WNT signaling (2-8), Receptor Tyrosine Kinase (RTK) signaling (9-16), Hippo signaling (17-22), and Jun N-terminal Kinase (JNK) signaling (23-25), as well as clear differences in regenerative mechanisms among organisms and tissues (26,27).

Assessing changes in gene expression in regenerating tissue is a powerful approach to identifying essential regeneration genes. Model organisms that are amenable to mutagenesis, transgenics, or RNAi-mediated gene knockdown also enable functional studies based on the results of transcriptional profiling. For example, analysis of the transcriptome of the cricket leg blastema identified upregulation of components of the Jak/STAT signaling pathway, which, when knocked down by RNAi, resulted in impaired leg regeneration (28). The transcriptome from the anterior of the planarian *Procotyla fluviatilis,* which is capable of regeneration after amputation, was compared to the transcriptome from posterior areas of the planarian body that are incapable of regeneration, identifying upregulation of several WNT ligands and receptors after amputation in the tissue that does not regenerate. RNAi knockdown of the WNT effector β-catenin restored regenerative capacity to the posterior of the animal (29). In zebrafish, genes regulating anterior-posterior patterning during fin regeneration were identified through transcriptional profiling of anterior and posterior portions of the blastema. Overexpression of one of these genes, *hand2* (SO:0000704), affected patterning but not growth during regeneration (30). Thus, transcriptional profiling followed by functional analysis is an effective approach to identification and validation of regeneration genes.

*Drosophila melanogaster* is one of the most powerful model organisms for genetic and functional analysis of genes. Furthermore, *Drosophila* imaginal discs, the epithelial structures in the larva that will form the adult animal during metamorphosis, have been an important model system for tissue repair and regeneration for over 60 years (reviewed in 31). This structure is a simple epithelium that contains complex patterning and determined cell fates. While classic imaginal disc regeneration experiments involved removal of the tissue from the larva before wounding and culturing in the abdomen of an adult host, the development of systems that use genetic tools to induce tissue ablation *in situ* has enabled high-throughput experimental approaches such as genetic screens (6,32). In both methods of inducing damage, the tissue undergoes wound closure and forms a regeneration blastema, or zone of proliferating cells near the wound (6,32-35). In addition, both methods of inducing damage activate signaling through the Wingless and JNK pathways (6,23,32,36-38).

Previous studies have identified genes differentially expressed during imaginal disc regeneration. Blanco et al. cut imaginal discs and then cultured them in the abdomens of adult female flies, before recovering the discs at various time points during regeneration for microarray analysis (39). This study used the entire imaginal disc for the microarrays, including tissue not contributing to the blastema. To restrict their analysis to cells near the wound site that were contributing to regeneration, Katsuyama et al. similarly cut and cultured discs, but used GFP-labeling of cells with activated JNK signaling to mark the regeneration blastema for dissection prior to microarray profiling (40). Together these studies used transcriptional profiling to identify several regeneration genes and mechanisms. However, they used relatively small numbers of cells from few regenerating discs due to the technical challenges inherent in the culture technique. Furthermore, culturing itself may induce high levels of stress in the tissue that may alter the transcriptional profile.

We sought to generate a complete and accurate transcriptional profile of regenerating imaginal disc tissue using deep-sequencing techniques and avoiding *ex vivo* culture and microdissections. Induction of tissue ablation using genetic tools enables regeneration to proceed *in vivo* as it would if the tissue were to be damaged by a predator or parasite in the wild. Furthermore, use of a genetic tissue-ablation system facilitates ablation and regeneration of hundreds of imaginal discs simultaneously, enabling collection of sufficient material for mRNA-seq without needing amplification. Finally, functional validation of the differentially expressed genes can be carried out by quantifying the extent and quality of regeneration after *in situ* tissue ablation in mutants.

Here we report the transcriptional profile of the regeneration blastema after ablation of the wing pouch in the *Drosophila* wing imaginal disc during the peak of regenerative growth. We have used a method that optimizes our ability to isolate fluorescently labeled blastema cells rapidly and efficiently from the disc (41), enabling collection of material for mRNA-seq. Furthermore, we have functionally validated several of the genes that are differentially expressed during regeneration as novel regulators of regeneration. We show that regulators of reactive oxygen species (ROS), in particular the DUOX-maturation factor NIP encoded by the gene *moladietz* (*mol*) (FBgn0086711) (42), play important roles in controlling ROS propagation in the regeneration blastema, which in turn helps sustain regeneration signaling and growth. The JNK signaling pathway, which is essential for regeneration (23) and is activated by ROS at the damage site (43), is also required for damage-induced upregulation of *mol*, demonstrating that a positive feedback loop between ROS production and JNK signaling sustains the regenerative response for several days after tissue damage. Thus, by whole-genome transcriptional profiling of regenerating tissue we have identified the changes in gene expression that control a key regulatory mechanism of regenerative growth.

## RESULTS

### Isolation of marked blastema cells

We induced ablation of most of the primordial wing by expressing the proapoptotic gene *reaper (rpr)* (FBgn0011706) (44) in the expression domain of the wing-patterning gene *rotund (rn)* (FBgn0267337) (45), which comprises most of the wing pouch region of the wing imaginal disc, via *rnGAL4, UASrpr* (6)(Fig 1A,B). To control the onset and completion of tissue ablation temporally, we used temperature shifts to regulate the temperature-sensitive repressor *Gal80^ts^* (46). We expressed *rpr* in the wing primordium for 24 hours at the beginning of the third larval instar, which removed most of the *rn*-expressing cells by the end of ablation to a reproducible extent (Recovery time 0 hrs or R0) (Fig 1 B)(Fig S1A). Wing pouch cells express the wing determinant *nubbin (nub)* (FBgn0085424)during both normal development and regeneration (6,47). Thus, *nub* expression was a convenient way to label blastema cells in these damaged discs as well as control cells in undamaged discs.

**Fig 1.**
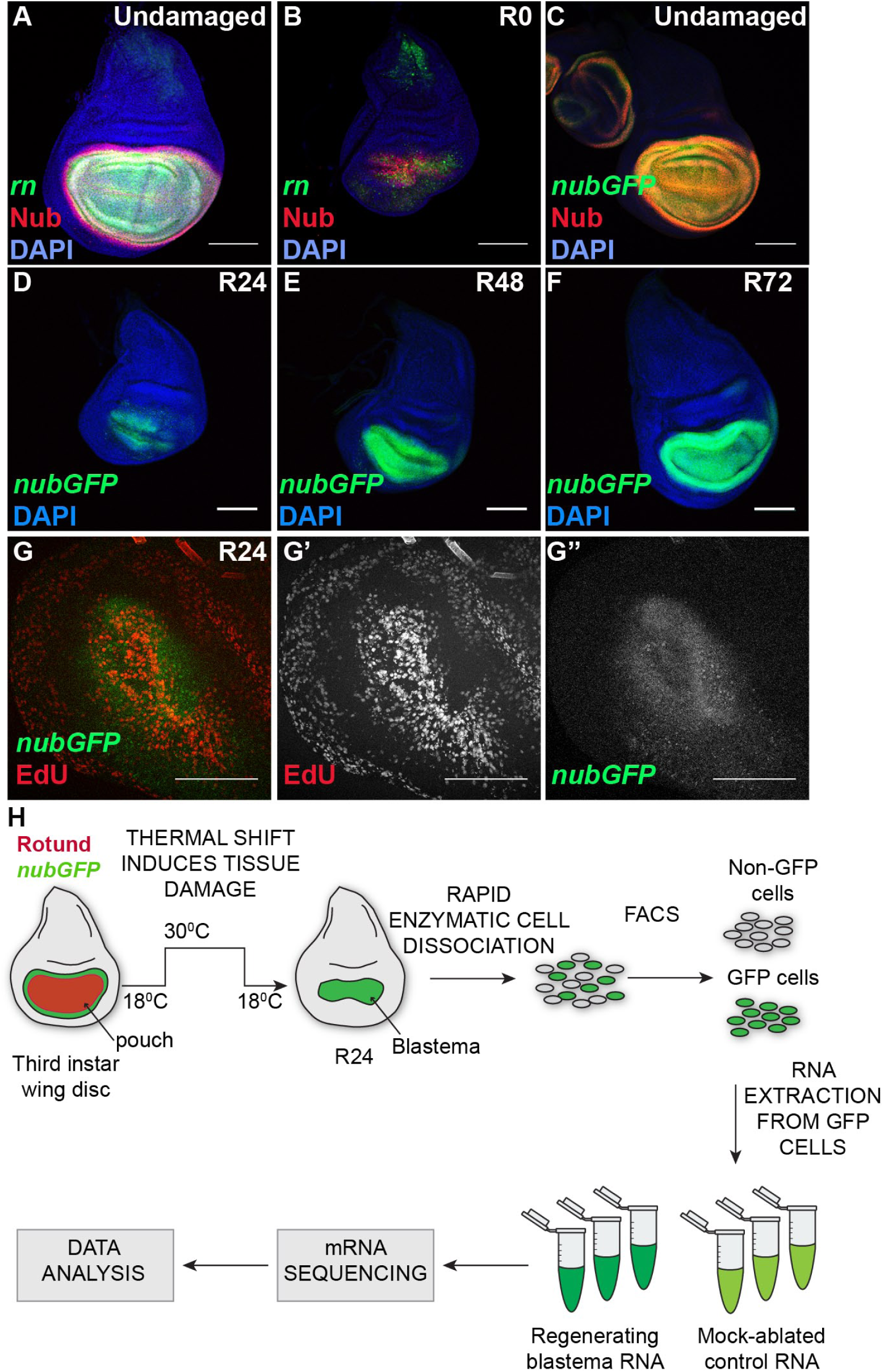
Labeling and isolating regeneration blastema cells. (A-B) Wing imaginal discs that are undamaged (A) or ablated at 0 hrs recovery (R0) (B). Green = *rnGal4, UAS-EGFP.* Red = anti-Nub. Blue = DAPI. (C) Wing imaginal disc showing overlap of anti-Nub immunostaining (red) and expression of the *nub-GFP* MiMIC enhancer trap (green). (D-F) *nub-GFP* marks the wing pouch at 24 hrs (D) 48 hrs (E) and 72 hrs (F) after ablation. (G) *nub-GFP* (green) coincides with the regeneration blastema as defined by a zone of high EdU incorporation (red). (H) Schematic of the mRNA-seq procedure, from tissue ablation through cell dissociation and sort to sequencing and data analysis. Scale bars are100 μm.

To label the regeneration blastema cells, we identified a publicly available MiMIC transposon insertion that expresses GFP under the control of the *nub* locus (48). Expression of GFP via this insertion occurs in the same cells that are immunostained with an anti-Nub antibody (Fig 1C) (41,47). The wing primordium continues to express the *nub-GFP* after ablation and throughout different stages of regeneration (Fig 1D-F). The GFP-expressing cells also encompass the regeneration blastema at R24 as marked by EdU incorporation (Fig 1G), confirming its suitability as a marker for blastema and control wing pouch cells.

To identify the differentially expressed genes in the blastema, we carried out transcriptional profiling of the GFP-labeled and isolated blastema cell population from R24 wing imaginal discs (Fig 1H). The R24 time point was chosen as it shows a clear blastema, whereas at earlier time points some discs had not yet formed the blastema, and at later time points some discs were beginning to re-pattern the regrown tissue. Dissociation and fluorescence–activated cell sorting (FACS) of imaginal disc cells is a well-established but lengthy procedure that may affect gene expression and cell viability (49,50). We therefore optimized our cell dissociation process so that it was rapid and gentle, taking approximately 15 minutes, to minimize changes in transcription and loss of cell viability due to the manipulation of the tissue (41). We have previously confirmed the accuracy of the sorting by using qPCR to measure expression of pouch and non-pouch genes in the sorted cells (41). After using this protocol to dissociate and sort regeneration blastema cells and control wing pouch cells, mRNA was prepared and pooled such that each biological replicate produced sufficient mRNA for deep sequencing (Fig 1H).

### Identifying differentially expressed genes in the blastema

To identify genes that are differentially expressed during imaginal disc regeneration, we collected three independent samples of *nub-GFP*-expressing blastema cells from regenerating discs and three independent samples of *nub-GFP*-expressing cells from undamaged ‘mock-ablated’ control discs after 24 hours of recovery from the thermal shift (R24). While the mock-ablated controls were taken through the thermal shift, they lacked *UAS-rpr* so did not ablate any tissue. Through deep sequencing we obtained approximately 27 million reads per replicate. Reads were aligned using Tophat2 (51,52) against the *Drosophila melanogaster* genome (NCBI, build 5.41). A total of 3,798 differentially expressed genes (p<0.05) were identified using Cuffdiff (51), with a false discovery rate of 0.5.

While a log2 fold change of 1.5 is often set as an arbitrary cutoff threshold for differentially expressed genes, our transcriptional profile showed a log2 fold change of 1.3 for the gene *puckered* (FBgn0243512), which is the phosphatase that is both a target and a negative regulator of JNK signaling in the regeneration blastema (23,53), prompting us to set our cutoff at 1.3. Thus, by selecting a cutoff of log2 fold change ≥ 1.3 or ≤ -1.3, p<0.05, we have identified 660 statistically significant differentially expressed genes, 504 of which are upregulated and 156 of which are downregulated in the regeneration blastema (Tables S1 and S2).

Several genes previously identified as imaginal disc regeneration genes were up-regulated in our transcriptional profile, including *dilp8* (54), *rgn* and *mmp1* (55), *puckered* (23), and *myc* (6). In addition, we found some overlap between our gene list and the differentially regulated genes noted in two previously reported transcriptional profiles of regenerating imaginal discs (Fig S2) (39,40). We compared the genes that were at least log2 1.3-fold up- or down-regulated in our dataset and those similarly at least 1.3-fold up-or down-regulated in the microarray analysis of posterior regenerating tissue 24 hours after damage presented in Katsuyama et al., in which they cut and cultured imaginal discs, and used GFP-labeling of cells with activated JNK signaling to mark the regeneration blastema for dissection prior to microarray profiling (40). There were 32 differentially expressed genes in common with this report (Fig S2). This profile led them to explore the role of JAK-STAT signaling in disc regeneration (40). Importantly, we also identified the JAK/STAT signaling ligand *upd/os* (FBgn0004956) as highly upregulated in the regenerating blastema. We also compared the genes that were at least log2 1.3-fold up- or down-regulated in our dataset and those listed as similarly up- or down-regulated in the figures and tables reporting the microarray analysis of whole regenerating discs 24 hours after damage presented in Blanco et al., as the whole list of 1,183 genes they identified as differentially expressed was not published (39). There were 10 differentially expressed genes in common with this report (Fig S2). For this analysis, they cut and cultured imaginal discs, and used whole discs for microarray profiling (39). The minimal overlap with previous studies may be due to several factors, including differences in method of wounding (cut vs. tissue ablation), the discs used (leg vs. wing), regeneration conditions (culture vs. in situ) and method of transcriptional profiling (microarray using few discs/blastemas vs. mRNA-seq using cells isolated from hundreds of blastemas). Furthermore, we have set a 1.3-fold threshold to define differentially expressed genes, and a reduction in this threshold would identify more overlap among these three studies. Strikingly, only one gene was upregulated in all three transcriptional profiles when using the 1.3-fold threshold: *yellow-*b (FBgn0032601), which is a target of JNK signaling during dorsal closure (56). Therefore, the current transcriptional profile will enable the study of previously unidentified regeneration genes and pathways.

To confirm that our transcriptional profile identified genes that were indeed differentially regulated in the imaginal disc blastema, we used antibodies, enhancer-trap lines, and protein-trap lines to visualize gene expression in undamaged and damaged wing discs. The “undamaged” control discs depict the expression of these genes during normal development. Of the 22 genes we tested, 16 (73%) were differentially expressed as predicted. Validated upregulated genes were *Alkaline phosphatase* 4 (*Alp4/Aph4*) (FBgn0016123) (57), *Atf3/A3-3* (FBgn0028550) (58), *chronologically inappropriate morphogenesis (chinmo)* (FBgn0086758) (59), *Ets21C* (FBgn0005660) (60), and *moladietz (mol)* (42)(Fig 2A-E). Other genes had expression patterns that changed from ubiquitous to restricted to the blastema, such as *fruitless (fru)* (FBgn0004652) (61), *LaminC* (FBgn0010397) (62), *AdoR* (FBgn0039747) (63), and *kayak (kay)* (FBgn0001297) (64) (Figs. 2F-G, S3A,B). A third class of genes showed strong upregulation around the blastema and slight upregulation in the blastema including *pickled eggs (pigs)* (FBgn0029881) (65) and a reporter for *Stat92E* (FBgn0016917) activity that reflects *upd*-stimulated signaling (66)(Fig 2H-I). The genes *Thor* (FBgn0261560) (67), *corto* (FBgn0010313) (68), *Nlaz* (FBgn0053126) (69), *twist (twi)* (FBgn0003900) (70), and *zfh1* (FBgn0004606) (71) showed upregulation in the transcriptional profile but did not show elevated expression with antibody staining (*twist*) or enhancer-trap expression (*zfh1, Thor* and *Nlaz*) or protein-trap expression (*corto)* (Fig S3C-G). Some of these genes may be upregulated in the transcriptional profile if, in the course of regeneration, hinge cells convert to pouch cells and begin expressing *nub* while still expressing some hinge-specific genes such as *zfh1*. Such hinge-to-pouch conversion has been reported during compensatory proliferation (72,73), and gene expression in these transitioning cells may still be important for regeneration.

**Fig 2.**
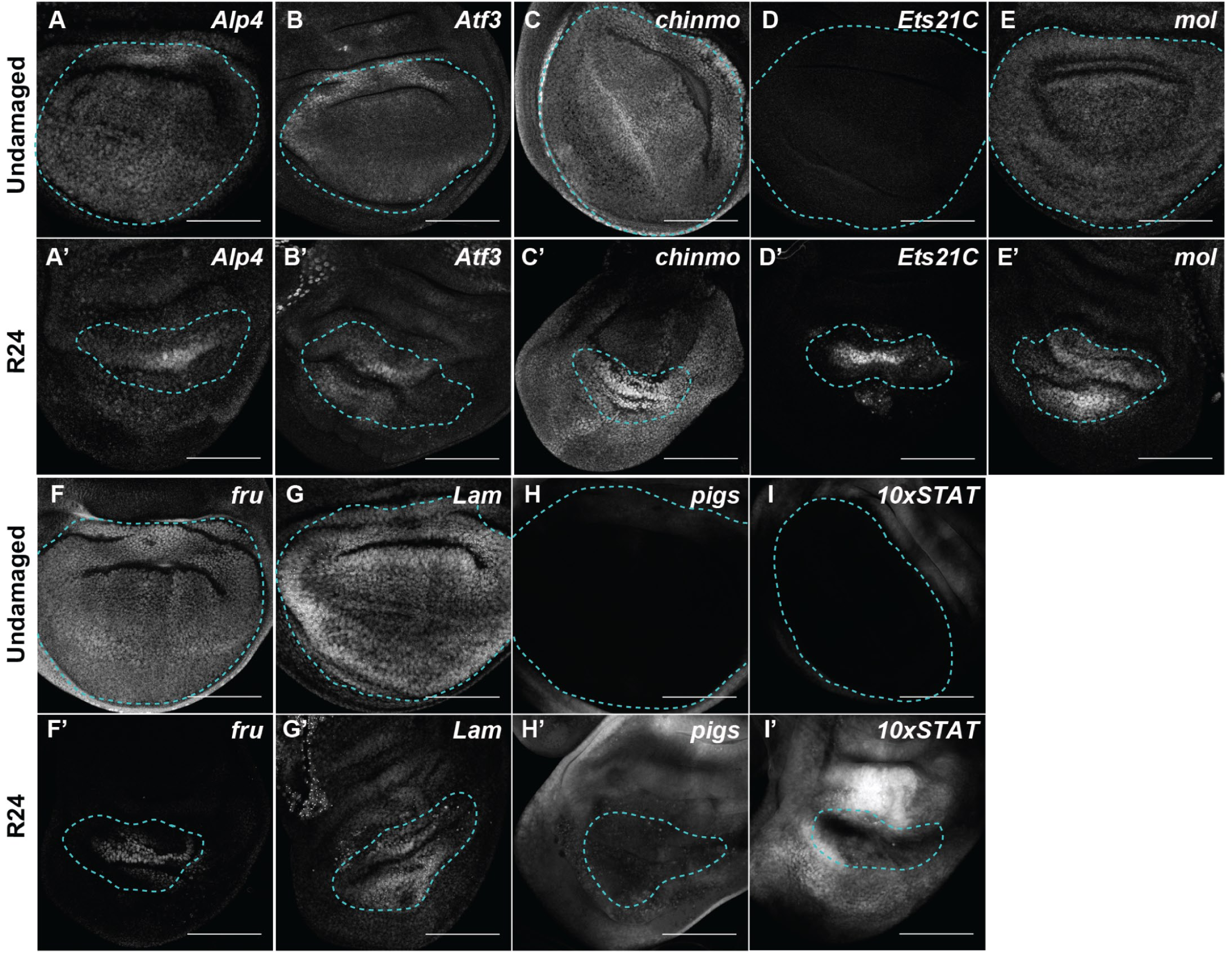
Validation of genes identified as upregulated in the regeneration blastema. Undamaged (A-I) and regenerating (A’-I’) wing discs. (A-A’) *Alp4-lacZ* enhancer trap. (B-B’) Atf3-GFP protein trap. (C-C’) *chinmo-lacZ* enhancer trap. (D-D’) Ets21C-GFP protein trap. (E-E’) *mol-lacZ* enhancer trap. (F-F’) *fru-lacZ* enhancer trap. (G-G’) Lamin-GFP protein trap. (H-H’) *pigs-GFP* enhancer trap. (II’) 10xSTAT92E-GFP reporter for STAT activity. Blue dashed line outlines the wing primordium. Scale bars are 100μm.

We also confirmed three of the upregulated genes using qPCR of whole wing discs (SFig 3H). While whole-disc qPCR often fails to detect differences in expression that occur only in the blastema, because the blastema consists of very few cells relative to the rest of the disc, changes in expression of genes that are largely not expressed in the disc prior to damage, such as *puckered,* can be observed (74). Thus, we further validated the upregulation of Ets21C, mol, and Nox as representatives of the differentially expressed genes (SFig 3H)

Validated downregulated genes were *defective proventriculus (dve)* (75), *Hormone receptor 78 (Hr78)* (FBgn0015239) (76), *NC2β* (FBgn0028926) (77), *smooth (sm)* (FBgn0003435) (78), *and Catalase (Cat)* (FBgn0000261) (79) (Fig 3). Thus, this transcriptional profile successfully identified genes that are differentially expressed in the regeneration blastema that forms after mass tissue ablation.

**Fig 3.**
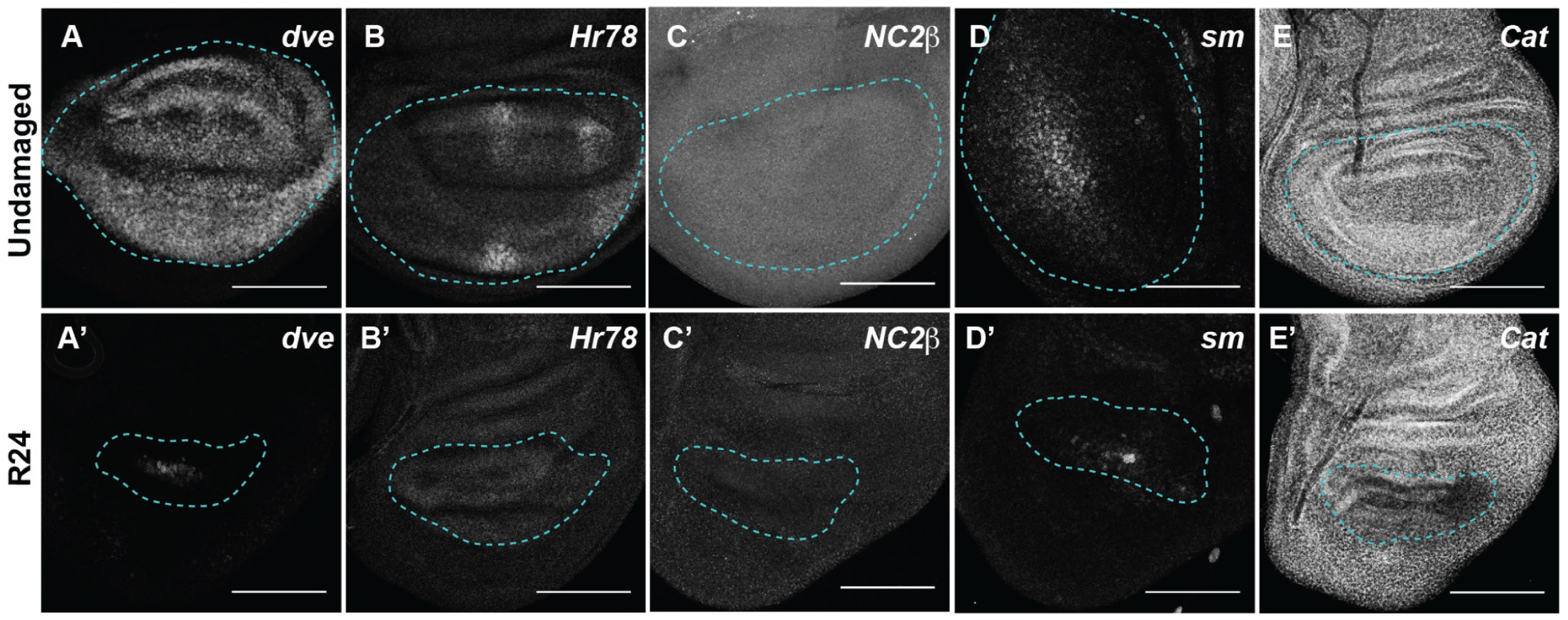
Validation of genes identified as downregulated in the regeneration blastema. Undamaged (A-E) and R24 (A’-E’) wing discs. (A-A’) *dve-lacZ* enhancer trap. (B-B’) Hr78-GFP protein trap. (C-C’) NC2β-GFP protein trap. (D-D’) *sm-lacZ* enhancer trap. (E-E’) *Cat-GFP* enhancer trap. Blue dashed lines outline the wing primordium. Scale bars are 100 μm.

### Identification of novel regeneration genes

A strong advantage to using a genetically tractable model organism is the ability to assess the functional role of genes of interest that are identified in a transcriptional profile. To assess regenerative capacity in the *Drosophila* imaginal wing, we induced tissue ablation as described above in animals that were heterozygous mutant for the gene in question. The regenerating animals were then allowed to develop to adulthood, and wing size was measured to assess the extent of regeneration. To measure a population of these wings efficiently, they were sorted into classes that were approximately <25%, 25%, 50%, 75%, and 100% the size of a normal wing (Fig 4A). The distribution of mutant regenerated wings in these classes was then compared to the distribution of regenerated wings generated by control animals. With our system, we observe some heterogeneity in the extent of regeneration within a genotype and also between control experiments conducted at different times. The variation within each genotype was due to variation in each individual animal’s time to pupariation, with animals that had longer to regenerate having larger wings (Fig S1B, C)(6). Variation between experiments was due to changes in environmental conditions such as humidity and food quality (74,80,81). Despite this apparent heterogeneity, we find reproducible differences between mutant and control animals using this method of screening and have successfully identified genes that regulate specific aspects of regeneration (6,74,81,82). Using this method, we tested available mutants in genes that were strongly upregulated after tissue damage. Twelve out of 16 or 75% of the genes we tested showed a regeneration phenotype, which is unsurprising given that not all important regeneration genes will have a phenotype when only heterozygous mutant, and not all differentially expressed genes will be essential for regeneration. One example of an upregulated gene that was required for regeneration is *Ets21c*, which encodes a transcription factor that is a known target of JNK signaling and is important for JNK activity in the innate immune response (83), during tumor formation (84,85) and at epidermal wounds (86). After ablation and regeneration of the imaginal tissue, adult wings in *Ets21c^f03639^*/+ animals were smaller than controls (Fig 4B). A second example of an upregulated gene that was required for regeneration is *CG9336* (FBgn0032897), which is annotated in the *Drosophila* genome and has closely related homologs in other *Drosophila* species but not in vertebrates, and does not appear to have protein domains of known function. After ablation and regeneration of the wing primordium in *CG9336^MI03849^*/+ animals, the resulting adult wings were smaller than controls, indicating a requirement for this gene during regeneration (Fig 4C). Additional genes required for regeneration included *alkaline phosphatase 4* (*Alp-4*) (57), the 4E-BP gene *Thor* (67), *moladietz (mol)* (42), as well as the collagen components *Collagen type IV alpha 1 (Col4a1/Cg25C)* (FBgn0000299)(87) and *viking (vkg)* (FBgn0016075)(88)(Fig 4D-G).

**Fig 4.**
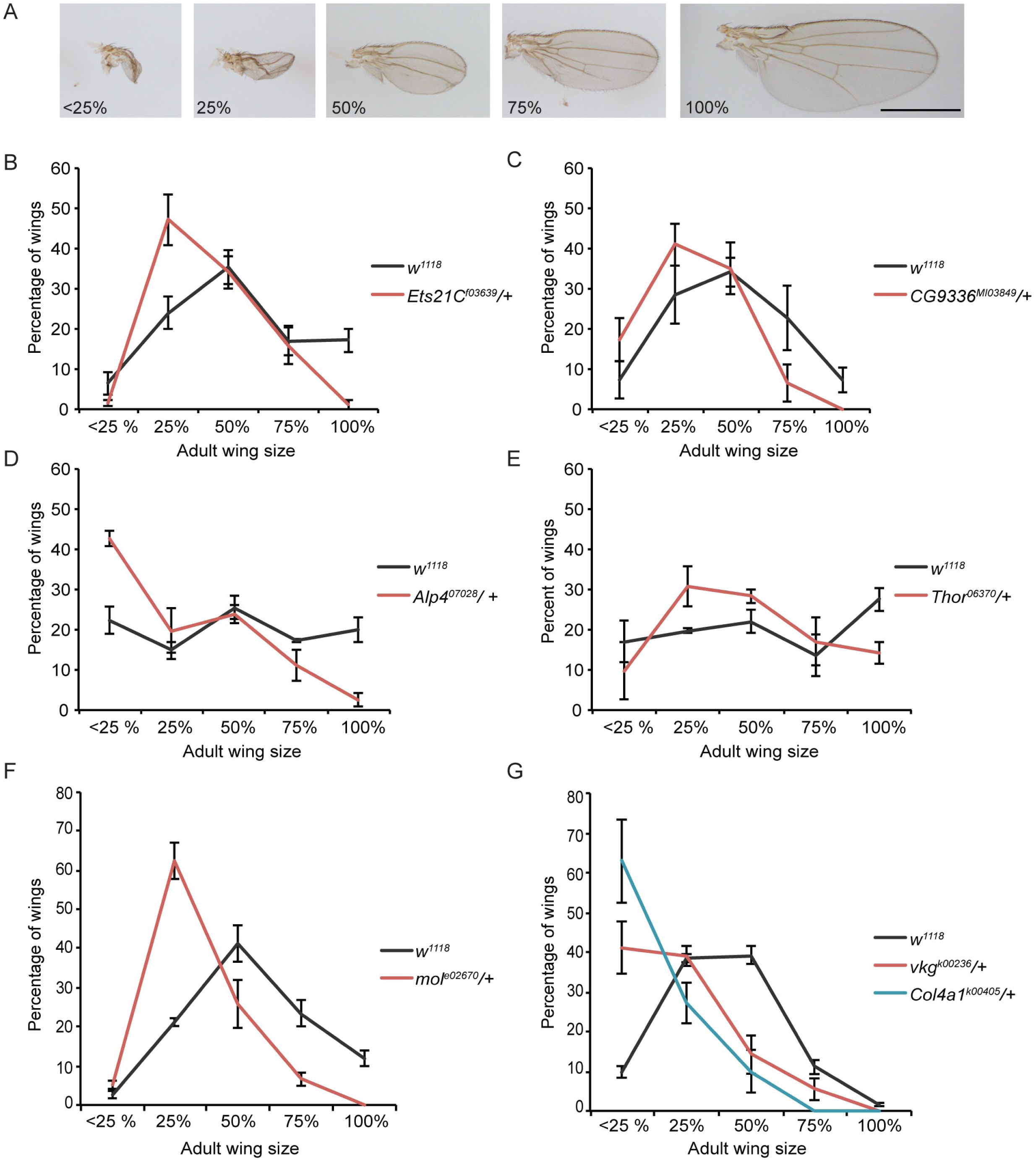
Genetic assays demonstrating that that differentially expressed genes have functional roles in regeneration. (A) Representative examples of wings from damaged discs that are approximately <25%, 25%, 50%, 75%, and 100% of a normal wing. Scale bar is 1mm. (B-G) Sizes of adult wings after regeneration in control (*w^1118^*) and heterozygous mutant animals. Three independent experiments each, error bars are SEM. (B) Sizes of adult wings after regeneration in *w^1118^* and *Ets21 C^f03639^*/+ animals. *w^1118^* n=318 wings, *Ets21C^f03639^*/+n=255 wings, p<0.0001 using a chi-squared test. (C) Sizes of adult wings after regeneration in *w^1118^* and *CG9336^MI03849^*/+ animals. *w^1118^* n=374 wings, *CG9336^MI03849^*/+ n=215 wings, p<0.0001 by a chi-squared test. (D) Sizes of adult wings after regeneration in *w^1118^* and *Alp4^07028^*/+ animals. *w^1118^* n=239 wings, *Alp4^07028^*/+ n=217 wings, p<0.0001 by a chi-squared test. (E) Sizes of adult wings after regeneration in *w^1118^* and *Thor^06270^*/+ animals. *w^1118^* n=224 wings, *Thor^06270^*/+ n=146 wings, p=0.0021 by a chi-squared test. (F) Sizes of adult wings after regeneration in *w^1118^* and *Mol^e02670^*/+ animals. Three independent experiments, *w^1118^* n=356 wings, *Mol^e02670^*/+ n=183 wings, p=0.00001 by a chi-squared test. (G) Sizes of adult wings after regeneration in *w^1118^*, *Col4a1^k00405^*/+, and *vkg^k00236^*/+ animals. *w^1118^* n=320 wings, *Col4a1^k00405^*/+ n=71 wings, and *vkg^k00236^*/+ n=134 wings, p<0.0001 by a chi-squared test.

Interestingly, several of the mutants tested did not have the predicted effect on regeneration. Rather than leading to poor regeneration, mutations in a subset of upregulated genes enhanced regeneration when heterozygous. These genes included *heartless (htl)* (FBgn0010389) (89), and *fru* (61) when assessed in males (Fig S4). After ablation and regeneration, wing sizes in these mutants were larger than control wings. The mechanisms through which these genes restrict regeneration are not yet understood.

### Biological processes affected during regeneration

To identify the biological processes that might be affected during regeneration we carried out gene ontology (GO) enrichment analysis. Transcripts that were significantly upregulated or downregulated were analyzed according to GO categories using DAVID v6.7 (90,91). Representative GO terms from the most significantly enriched GO clusters describing biological processes are listed in Table 1. Terms that were enriched among the upregulated genes included imaginal disc development and imaginal disc pattern formation, likely because the regenerating tissue was rebuilding what had been ablated. The enrichment of GO terms cell morphogenesis, tissue morphogenesis, cell adhesion, morphogenesis of an epithelium, and cell migration may occur because the cells at the wound edge change shape in order to close the wound (32). In addition, discontinuity in marked clones in regenerating tissue suggests that cells intercalate and shift relative to each other during imaginal disc regeneration (92). The GO term regulation of transcription likely contains transcription factors necessary for carrying out the regeneration program,as well as the development and patterning of the regenerating tissue. Interestingly, the GO term open tracheal system development was highly enriched. Two possible reasons for this apparent enrichment include contamination of our sorted cells with tracheal cells, or upregulation in the blastema of the same RTK signaling pathway genes that play critical roles in tracheal system morphogenesis, as has been observed in compensatory proliferation (93). Another highly enriched GO cluster included the terms negative regulation of cell differentiation, regulation of cell fate commitment, and regulation of cell fate specification. Interestingly, we and others have shown that imaginal disc damage causes a transient loss of markers of cell-fate specification (6,94).

**Table 1:**
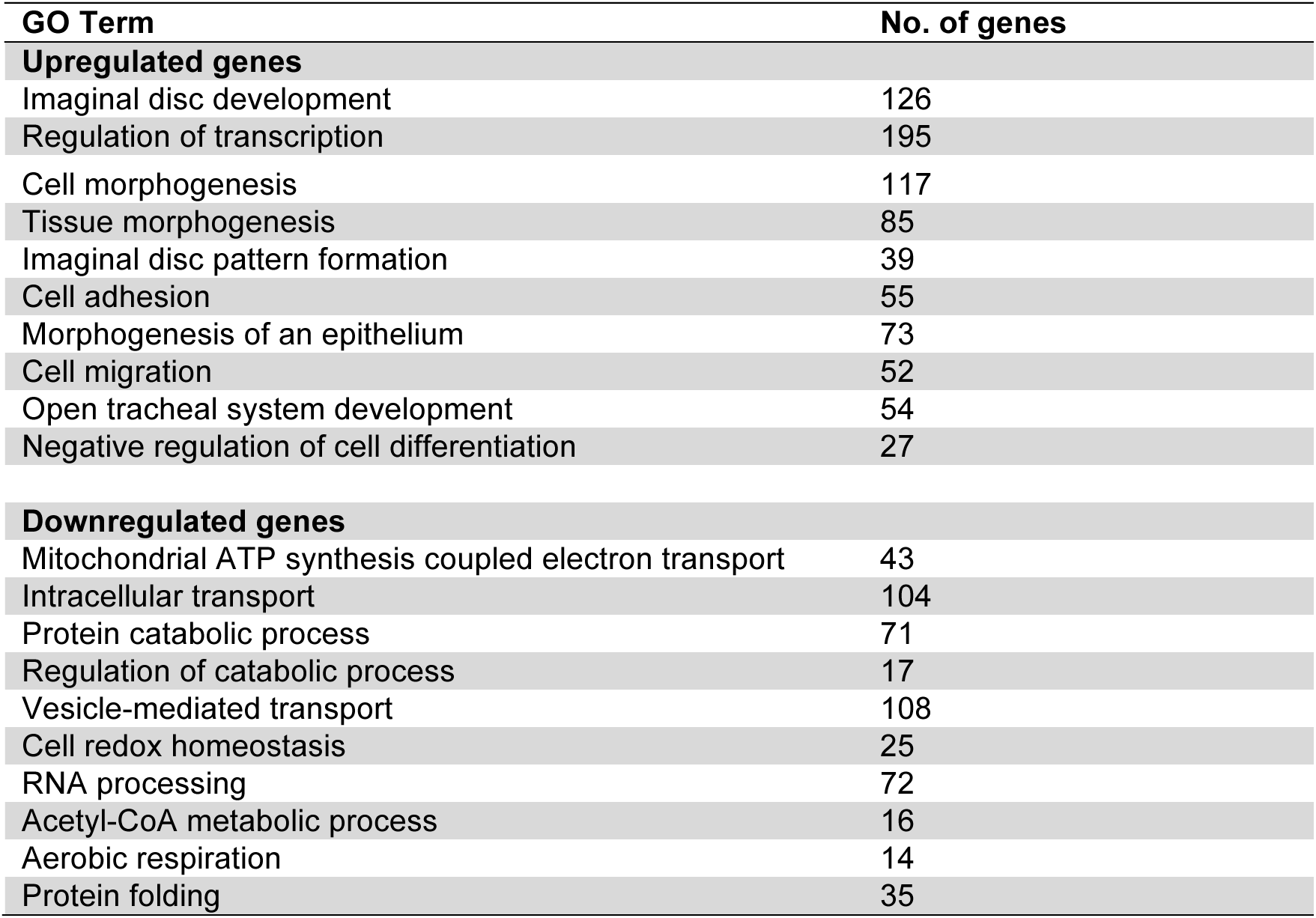
Gene ontology (GO) analysis for enrichment of biological process terms

| GO Term | No. of genes |
| --- | --- |
| **Upregulated genes** |  |
| Imaginal disc development | 126 |
| Regulation of transcription | 195 |
| Cell morphogenesis | 117 |
| Tissue morphogenesis | 85 |
| Imaginal disc pattern formation | 39 |
| Cell adhesion | 55 |
| Morphogenesis of an epithelium | 73 |
| Cell migration | 52 |
| Open tracheal system development | 54 |
| Negative regulation of cell differentiation | 27 |
| **Downregulated genes** |  |
| Mitochondrial ATP synthesis coupled electron transport | 43 |
| Intracellular transport | 104 |
| Protein catabolic process | 71 |
| Regulation of catabolic process | 17 |
| Vesicle-mediated transport | 108 |
| Cell redox homeostasis | 25 |
| RNA processing | 72 |
| Acetyl-CoA metabolic process | 16 |
| Aerobic respiration | 14 |
| Protein folding | 35 |

Many of the biological process GO terms enriched among the downregulated genes describe general cellular processes, including intracellular transport and vesicle-mediated transport, RNA processing, and catabolism. Interestingly, several classes of genes that affect cellular metabolism were downregulated, including the GO terms mitochondrial ATP synthesis coupled electron transport, acetyl-A metabolic process, tricarboxylic acid cycle, and aerobic respiration. While a transcriptional profile of the *Xenopus tropicalis* tadpole tail regenerative bud has similarly suggested changes in cellular metabolism after tissue damage (95), a broad, functional role for cell-autonomous changes in oxidative phosphorylation, glycolysis or other cellular energetics during regeneration has yet to be demonstrated.

### Regulators of Reactive Oxygen Species are differentially expressed during regeneration

An additional downregulated GO category was cell redox homeostasis, suggesting changes in levels of enzymes that regulate Reactive Oxygen Species (ROS) in the regeneration blastema. Indeed, ROS provide important signaling in other model systems of wound healing and regeneration (reviewed in 96). For example, ROS serve as an attractant for immune cells in larval zebrafish tails after amputation (97) and in *Drosophila* cuticle wounds (98), and are required for proliferation and regeneration after *Xenopus* tadpole tail amputation (99) as well as fin and axon regrowth after zebrafish tail amputation (100,101). Furthermore, ROS stimulate JNK signaling in regenerating zebrafish fins and *Drosophila* imaginal discs (43,102,103). During wing imaginal disc regeneration ROS are released by the dying cells, and then taken up by the living cells at the wound edge immediately after physical damage or induction of tissue ablation (43). However, the extent to which ROS are produced and propagated in the regeneration blas-tema, as well as the mechanism that underlies ROS production in the regrowing tissue, are unclear.

We examined the expression of genes that regulate ROS production and removal in our transcriptional profile of the imaginal disc regeneration blastema, and found that in addition to the downregulated genes identified by the GO analysis, there were also ROS-regulating factors among the upregulated genes (Table 2). *Drosophila* has two NADPH oxidases that produce ROS, NADPH Oxidase (Nox) (FBgn0085428) and Dual oxidase (Duox) (FBgn0283531) (104-107). Interestingly, *Nox* expression was upregulated, while *Duox* expression remained unchanged. However, the Duox-maturation factor DUOXA/NIP, which is encoded by the gene *moladietz (mol)* (42), showed a high level of induction after damage, representing one of the strongest hits in the profile. To reduce ROS, superoxide and hydrogen peroxide are scavenged by superoxide dismutases (Sods) and Catalase (Cat), respectively. Expression of the CuZn-dependent cytoplasmic *Sod1* (FBgn0003462)(108) and the Mn-dependent mitochondrial *Sod2* (FBgn0010213)(109) was reduced in the regeneration blastema, while the extra-cellular *Sod3* (FBgn0033631)(110) remained unchanged. Furthermore, expression of *Cat* (79) was strongly reduced. Thus, generation and propagation of ROS in the regeneration blastema could be explained in part by transcriptional upregulation of Nox and *mol*/NIP, and downregulation of Sod1, Sod2, and Cat.

**Table 2:**
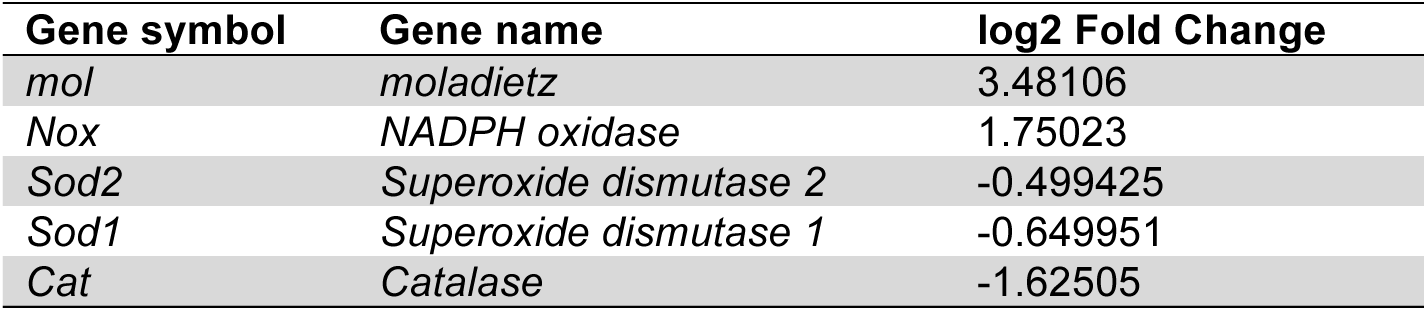
List of reactive oxygen species (ROS) regulating genes

| Gene symbol | Gene name | log2 Fold Change |
| --- | --- | --- |
| *mol* | *moladietz* | 3.48106 |
| *Nox* | *NADPH oxidase* | 1.75023 |
| *Sod2* | *Superoxide dismutase 2* | -0.499425 |
| *Sod1* | *Superoxide dismutase 1* | -0.649951 |
| *Cat* | *Catalase* | -1.62505 |

### ROS is required to sustain regeneration signaling

While regenerating zebrafish tails exhibit ROS production for at least 24 hours after amputation (102), and Xenopus tadpole tails produce ROS for days after amputation (99), ROS production in damaged wing discs has only been assessed for 30 minutes after physical damage and 11 hours after induction of tissue ablation (43). To determine whethROS-regulating enzymes impact regenerationer ROS persist in regenerating wing discs, we used dihydroethidium (DHE) staining to detect ROS. Importantly, we observed DHE fluorescence in the cellular debris and in the regeneration blastema at R24 (Fig 5A-B) and R48 (Fig 6F). We confirmed this finding with the ROS detector H_2_DCFDA (SFig 6D,E). Thus, ROS persist in the living, regenerating cells for at least 24 hours after the completion of tissue ablation, suggesting an active mechanism that sustains the production of ROS in the regenerating tissue.

**Fig 5.**
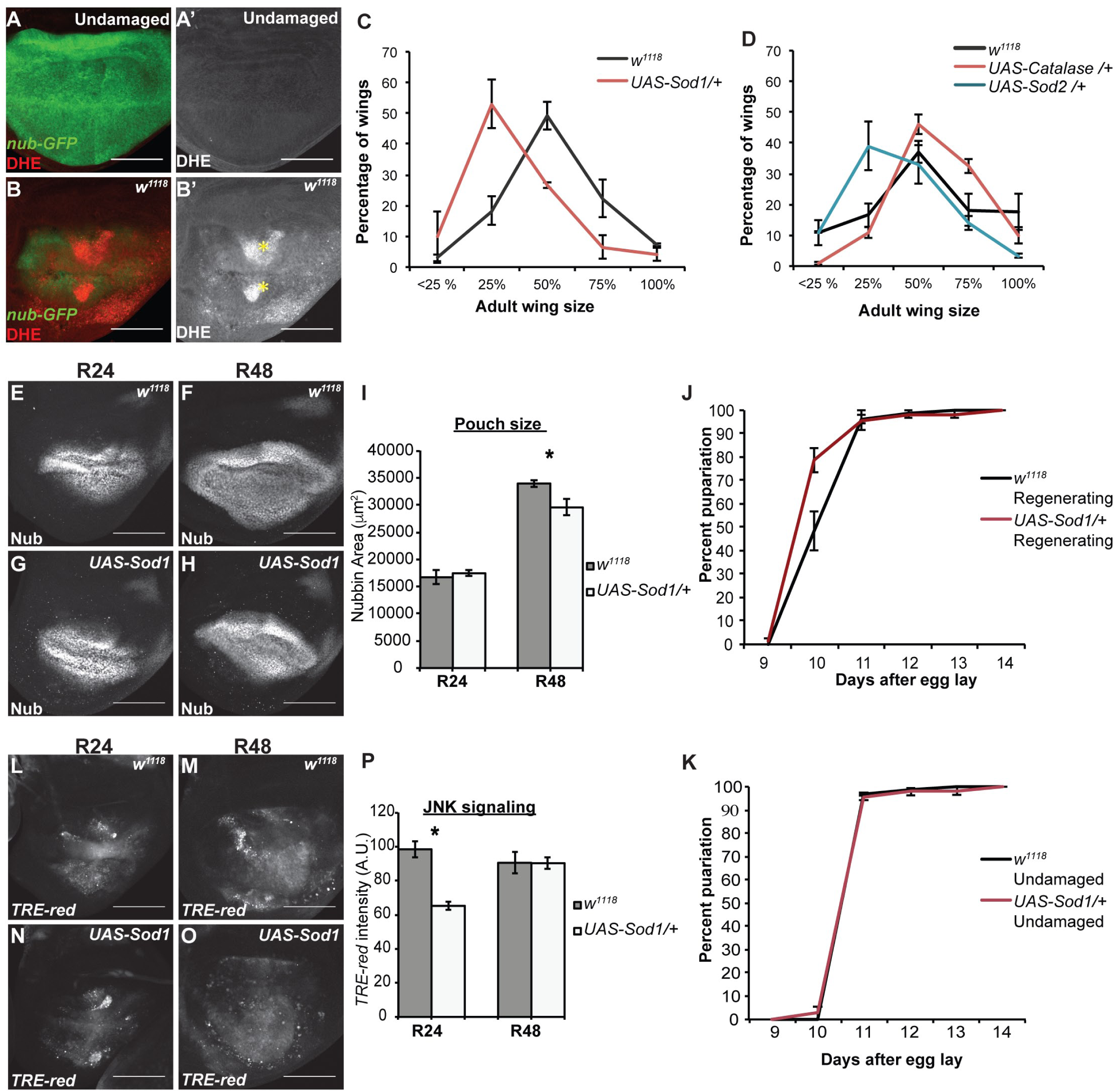
ROS persist in the regeneration blastema and are required for regeneration. (A-B) DHE staining (red) to detect ROS. The wing pouch is marked with *nub-GFP* (green). Yellow asterisks mark pockets of cellular debris. (A) Undamaged disc. (B) Regenerating disc at R24. (C-D) Genetic regeneration assays using adult wing size to assess extent of regenerative growth in the imaginal discs. Three independent experiments for each. (C) Sizes of adult wings after regeneration in *w^1118^* and *UAS-Sod1*/+ animals. *w^1118^* n=375 wings, *UAS-Sod1*/+ n=166 wings, p<0.0001 using a chi-squared test. (D) Sizes of adult wings after regeneration in *w^1118^*, *UAS-Sod2*/+, and *UAS-Cat*/+ animals. *w^1118^* n=327 wings, *UAS-Sod2*/+ n=332 wings, *UAS-Cat*/+ n=361 wings, p<0.0001 using a chi-squared test. (E-H) Anti-Nub marks the wing primordium of *w^1118^* (E,F) and *UAS-Sod1* (G,H) regenerating discs at R24 and R48. (I) Quantification of area of the wing primordium as marked by anti-Nub at R24 and R48. *w^1118^* R24 total n=12 discs, *UAS-Sod1* R24 n=15 discs, *w^1118^* R48 n=5 discs, *UAS-Sod1* R48 n=10 discs. At R48, p=.0248. (J-K) Pupariation rates. Note that because of the temperature shifts in the ablation protocol the regenerating and undamaged pupariation times cannot be compared to each other. (J) Pupariation timing after regeneration. Three replicates, control n=213 pupae, *UAS-Sod1* n=107 pupae. (K) Pupariation timing during normal development. Three replicates, control n=173 pupae, *UAS-Sod1* n = 201 pupae. (L-O) Expression of the *TRE-red* reporter indicates JNK signaling activity in *w^1118^* (L,M) and *UAS-Sod1* (N,O) regenerating discs at R24 and R48. (P) Quantification of *TRE-red* fluorescence as an indicator of level of JNK signaling at R24 and R48. *p<.00002. *w^1118^* R24 n=10 discs, *UAS-Sod1* R24 n=14 discs, *w^1118^* R48 n=10 discs, *UAS-Sod1* R48 n=14 discs.Scale bars are 100 μm. Error bars are SEM.

**Fig 6.**
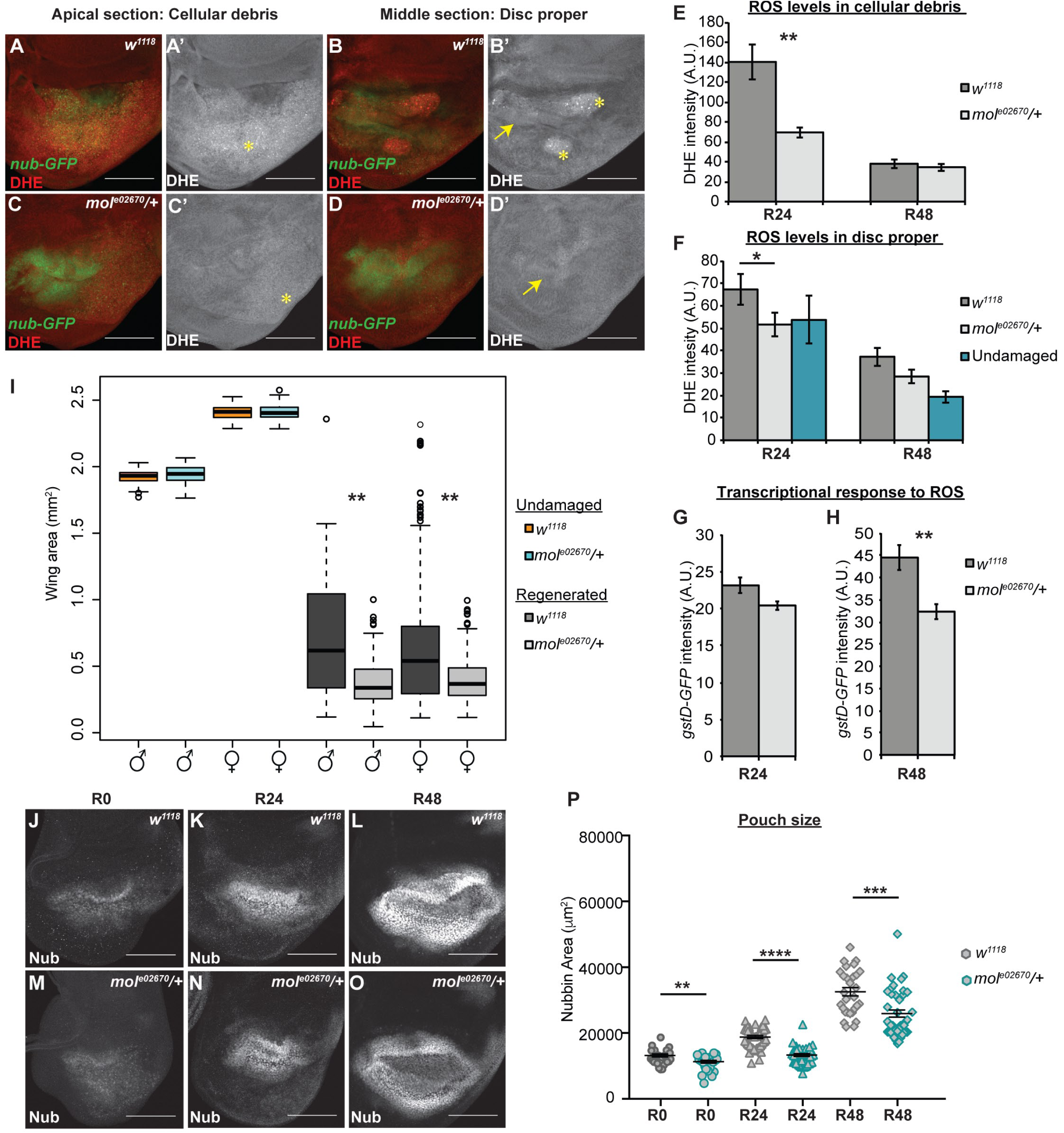
*moladietz* is required for wing disc regeneration. (A-D) DHE fluorescence (red) indicates the presence of ROS. *nub-GFP* (green) marks the regenerating wing pouch. (A-B) Confocal slices of a *w^1118^* regenerating disc through the debris field (A,A’) and the disc epithelium (B,B’). Asterisks mark cellular debris in the debris field and in a few folds in the epithelium. Arrow points to the position of the regenerating wing pouch. (C-D) Confocal slices of a *mol^e02670^*/+ regenerating disc through the debris field (C,C’) and the regenerating epithelium (D,D’). Asterisk and arrow same as above. (E-F) Quantification of DHE fluorescence intensity in the debris fields of *w^1118^* and *mol^e02670^*/+ regenerating discs (E) and in the regenerating epithelia of *w^1118^* and *mol^e02670^*/+ regenerating discs and control undamaged discs (F). For R24, three independent experiments, *w^1118^* regenerating n=12 discs, *mol^e02670^*/+ regenerating n=18 discs, *w^1118^* undamaged n=11 discs. For R48, three independent experiments for a total *w^1118^* regenerating n=30 discs, *mol^e02670^*/+ regenerating n=25 discs,*w^1118^* undamaged n=10 discs. (G,H) Quantification of GFP fluorescence from a *gstD-GFP* reporter for ROS-regulated transcription in regenerating *w^1118^* and *mol^e02670^*/+ discs. For R24, *w^1118^* n=12 discs, *mol^e02670^*/+ n=20 discs. For R48, *w^1118^* n=12 discs, *mol^e02670^*/+ n=10 discs. (I) Adult wing area in *w^1118^* and *mol^e02670^*/+ male and female wings from undamaged discs and after disc regeneration. Three independent experiments. Undamaged: *w^1118^* females n=125 wings, *w^1118^* males n=132 wings, *mol^e02670^*/+ females n= 82 wings, *mol^e02670^*/+ males n=73 wings. Regenerated: *w^1118^* females n=226 wings, *w^1118^* males n=134 wings, *mol^e02670^*/+ females n= 128 wings, *mol^e02670^*/+ males n=133 wings. (J-O) Anti-Nub marks the regenerating wing primordium at R0, R24 and R48 in *w^1118^* and *mol^e02670^*/+ discs. (P) Quantification of the size of the regenerating wing primordium at R0, R24 and R48. R0 *w^1118^* n=26 and *mol^e02670^*/+ n=29, R24 *w^1118^* n=42 and *mol^e02670^*/+ n=41, R48 *w^1118^* n=29 and *mol^e02670^*/+ n=42. Scale bars are 100 μm. Error bars are SEM. **p<0.05, *p<0.005, ***p<0.0002, ****p<0.0001

To determine the extent to which changes in ROS levels impact regeneration, we overexpressed Sod1, Sod2, or Cat in ablated discs using a *UAS-Sod1* (111), *UAS-Sod2* (112), or *UAS-Cat* (113) transgene under the control of *rn-GAL4*, which induced expression in the ablated tissue as well as in the few surviving *rn*expressing cells that contributed to the blastema. This limited overexpression was intended to reduce ROS levels in the debris and partially reduce ROS levels in the blastema, as not all blastema cells expressed the transgenes. According to a prior report, similar overexpression of Sod1 or Cat individually or together reduced the ability of wing discs to recover from tissue ablation (43). In our ablation system, overexpression of Sod1 or Sod2 during ablation similarly led to smaller adult wings compared to controls, although overexpressing Cat alone did not, confirming that manipulation of levels of ROS-regulating enzymes impact regeneration (Fig 5C,D). This reduction in regeneration was likely due to a combination of reduced regenerative growth, as *UAS-Sod1* regenerating wing primordia lagged behind controls in size (Fig 5 E-I), and reduced time for regeneration, as *UAS-Sod1* regenerating animals failed to delay pupariation in response to the tissue damage as long as the controls (Fig 5J,K). The damaged discs with transiently overexpressed Sod1 also had reduced JNK signaling at R24, as observed by expression of the *TRE-red* transcriptional reporter for JNK pathway activity (114) (Fig 5 L-P).

In damaged eye imaginal discs, ROS recruit hemocytes to the site of damage, which then stimulate JNK signaling in the recovering epithelium (103). To determine whether hemocytes are recruited to the wing disc in response to ablation of the *rn*-expressing domain, we observed hemocytes using Hemolectin-RFP (FBgn0029167) (115) and anti-Nimrod (FBgn0259896) (116). In control discs, small clusters of hemocytes were observed in the folds of 1 of 15 discs (Fig S5AB). In damaged and regenerating tissue, clusters of hemocytes were observed in 4 of 15 discs along the peripodial epithelium. These hemocytes were present in close proximity to the debris, but were not in direct contact with the debris, which was trapped between the two epithelial layers (Fig S5C-G). Thus, in contrast to the eye disc, recruiting hemocytes is unlikely to be the main mechanism through which ROS induce JNK signaling in the wing disc.

### The DUOX maturation factor *moladietz/*NIP is required for ROS production in the regeneration blastema

The striking upregulation of *mol* in the regeneration blastema by R24 and its continued expression through R48 (Fig 2E, SFig 8A-E) suggested that its protein product NIP may have an important role in regulating regeneration. Importantly, *mol* is normally expressed at low levels in the wing disc during development (Fig 2E, SFig 6A,E). The vertebrate homolog of NIP, DUOX maturation factor (DUOXA) (HGNC:26507), is essential for moving DUOX through the endoplasmic reticulum and the golgi to the cell surface (117). Once at the cell surface, DUOXA remains in a stable complex with DUOX and enhances the rate and specificity of ROS production (118). Thus, transcriptional regulation of *mol* could have a profound effect on ROS production in the regenerating epithelium.

To determine the extent to which the transcriptional upregulation of *mol* promotes ROS production in the blastema, we assessed ROS levels in heterozygous *mol* null mutant animals. Duox can produce both hydrogen peroxide and superoxide (118). As DHE was the reagent that worked best in imaginal discs (Fig 5A,B, SFig 6D,E), we used it as a representative assay for overall ROS levels. Interestingly, production of ROS in both the cellular debris and the regeneration blastema was significantly reduced in the *mol^e02670^*/+ damaged discs (Fig 6A-F), indicating that *mol* is required for overall ROS production after tissue damage. To confirm that the response to ROS is reduced in the *mol^e02670^*/+ regenerating tissue, we assessed expression of a reporter transgene, *gstD1-GFP* (FBgn0001149), that responds to ROS-induced activation of transcription (119). Interestingly, *gstD1-GFP* expression was significantly reduced in the mutant regeneration blastemas, but not until two days after tissue damage (Fig 6 G,H; SFig 6 F-J).

Our initial genetic assay showed that NIP was required for regeneration (Fig 4F). To quantify the effect of reduction of NIP further, we measured adult wing size in *mol^e02670^*/+ females and males after imaginal disc ablation and regeneration. Importantly, while normal wings were the same size in controls and *mol^e02670^*/+ animals, regenerated *mol^e02670^*/+ wings were significantly smaller than regenerated controls, indicating that regeneration in these *mol^e02670^*/+ animals was impaired (Fig 6I). To confirmed the requirement for *mol* we also quantified regeneration using discs expressing a *UAS-molRNAi* in the regenerating wing pouch. While such RNAi expression was limited temporally and spatially, we have found it to be effective at generating phenotypes in our system (74), possibly due to the propagation of knockdown after limited RNAi expression observed in imaginal discs (120). Importantly, discs *UAS-molRNAi* also regenerated worse than controls as assessed by adult wing size (SFig 6A).

To understand how reduced expression of *mol* impairs regeneration, we monitored regrowth of the ablated tissue by measuring the area of the wing primordium at specific times after the completion of ablation. We found that *mol^e02670^*/+ regenerating discs were slightly smaller than controls beginning in early regeneration, and significantly lagged behind controls in size by two days after tissue damage (Fig 6J-P).

### The DUOX maturation factor *moladietz/*NIP is required for sustained JNK signaling during regeneration

Given that the difference in regrowth was more apparent later in regeneration, we speculated that reduction of NIP levels might be particularly important for the later stages of regeneration. Indeed, expression of the growth-promoter Myc (FBgn0262656), which is important for regenerative growth (6) was comparable to controls at R24 but reduced at R48 (Fig S6K-O). Because ROS stimulate JNK signaling in damaged imaginal discs (43), we examined JNK signaling levels in *mol^e02670^*/+ regenerating discs. Importantly, expression of the JNK signaling reporter *TRE-red*, which reflects the activity of the AP-1 transcriptional complex, was slightly reduced during early and mid regeneration (R0, R24, and R48) and markedly reduced during the late stages of regeneration in *mol^e02670^*/+ discs (R72) (Fig 7A-J). To determine whether increasing JNK signaling could compensate for the reduction of NIP levels and ROS production, we examined adult wings after damage and regeneration in animals heterozygous mutant for both *mol* and the negative regulator of JNK signaling *puckered* (*puc*) (53). These *mol^e02670^/+; puc^E69^*/+ regenerated wings were significantly larger than the *mol^e02670^*/+ regenerated wings, indicating that increased JNK signaling could bypass the requirement for *mol* and rescue the poor regeneration phenotype of the *mol^e02670^*/+ mutants (Fig 7K). Thus, upregulation of *mol* is required for ROS propagation in the regeneration blastema and for sustaining JNK signaling, particularly during the later stages of regeneration.

**Fig 7.**
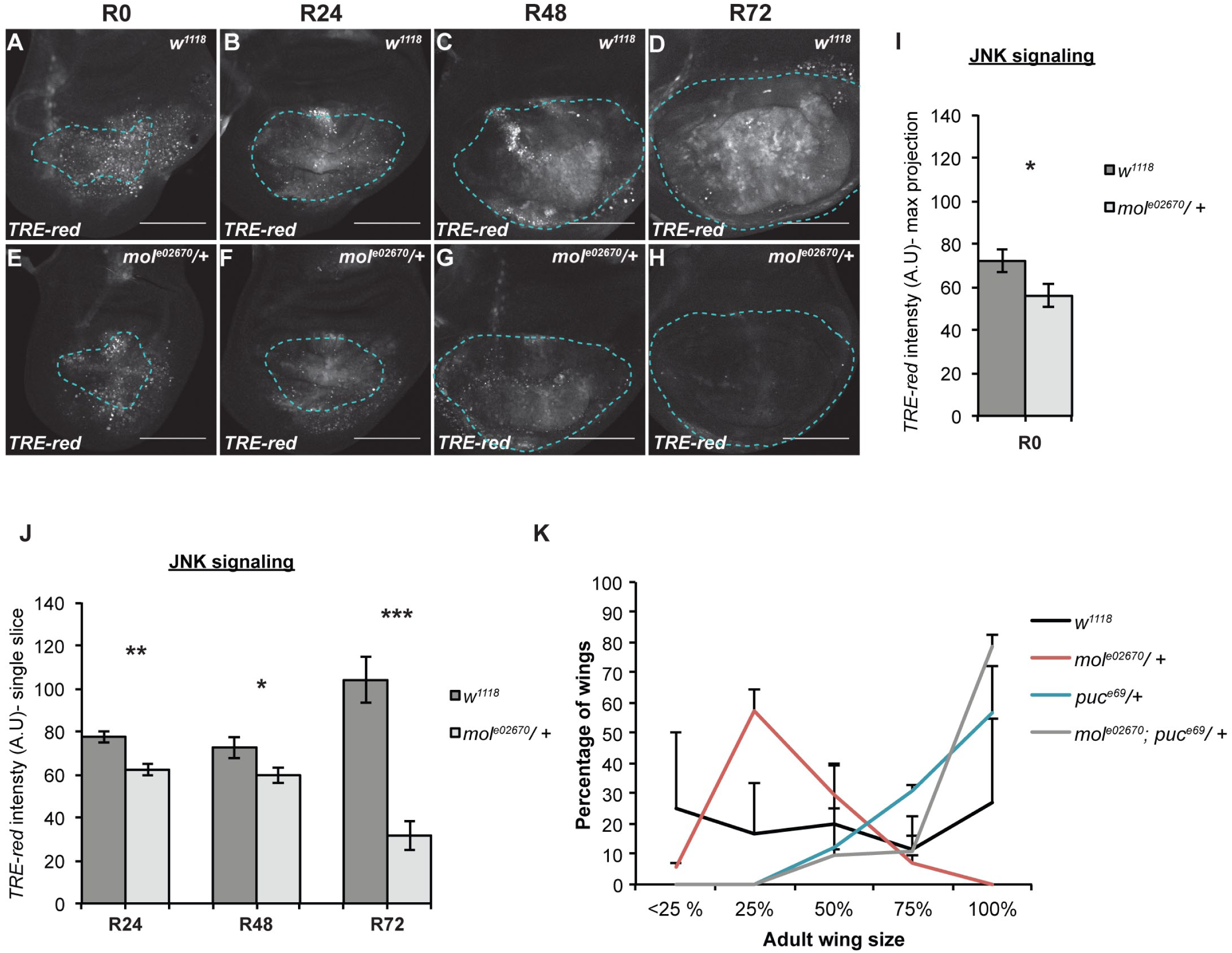
NIP is required to sustain JNK signaling during late regeneration. (AH) Confocal images of fluorescence from the *TRE-red* reporter for JNK signaling in *w^1118^* (A-D) and *mol^e02670^*/+ (E-H) regenerating discs at R0 (A,B), R24 (B,F), R48 (C,G) and R72 (D,H). (I) Quantification of fluorescence intensity of the *TREred* reporter in max projections of the confocal images at R0, because at this time point the epithelium cannot be distinguished from the debris.*w^1118^* n=10 discs, *mol^e02670^*/+ n=14 discs. (J) Quantification of fluorescence intensity of the *TRE-red* reporter in single slices of the confocal images through the regenerating epithelium at R24, R48, and R72. R24 *w^1118^* n=11 discs, *mol^e02670^*/+ n=11 discs. R48 *w^1118^* = 14 discs, *mol^e02670^*/+ n=15 discs. R72 *w^1118^* n=11 discs, *mol^e02670^*/+ n=11 discs. (K) Regeneration assays using adult wing size to assess extent of regenerative growth in the imaginal discs in *w^1118^*, *mol^e02670^*/+, *puc^E69^/+,* and *mol^e02670^/+;puc^E69^*/+ animals. Two independent experiments, thus error bars are SD. *w^1118^* n=26 wings, *mol^e02670^*/+ n=83 wings, *puc^E69^*/+ n=99 wings, and *mol^e02670^/+;puc^E69^*/+ n=95 wings. p<0.0001 for all comparisons using a chi-squared test. Dashed blue line outlines the wing primordium. Scale bars are 100 μm. Error bars are SEM unless otherwise noted. *p<0.05, **p<0.001, ***p<0.0001

The importance of the Duox-maturation factor in regeneration implies that Duox itself is also important for regeneration, even though it is not transcriptionally up-regulated according to our profile. To assess the importance of Duox, we quantified regeneration in *UAS*-*duoxRNAi* animals. Indeed, wing discs expressing *duoxRNAi* regenerated poorly compared to control animals (Fig S7 A,D).

Another important regulator of ROS that was upregulated in the transcription profile is the NADPH oxidase Nox. To determine whether Nox is also required for wing disc regeneration, we compared adult wings after damage and regeneration in control animals and animals heterozygous for a *Nox* mutant (*Nox^MI15634^*) or expressing *NoxRNAi*. Interestingly, both the *Nox* mutation and the *NoxRNAi* caused improved regeneration as assessed by adult wing size (Fig S7 B,C,E). To understand why reduction of Nox led to enhanced regeneration, we assessed pouch size throughout regeneration and rate of pupariation, Interestingly, wing discs with reduced Nox regrew at the same rate at control discs through R48, and pupariation timing was not altered (Fig S7 F-I). Thus, the constraint Nox places on regeneration must occur after R48, possibly during the pupal phase. These results suggest that the ROS produced by Nox and by the Duox/NIP complex are likely functionally, spatially, or temporally different, with Nox-produced ROS acting to inhibit regeneration during the pupal phase.

### JNK signaling is required for the upregulation of *mol* expression after tissue damage

Given that *mol* upregulation after tissue damage was important for ROS production in the regenerating epithelium and sustained regenerative signaling, we wanted to identify the upstream signal that regulates *mol* expression. We hypothesized that regeneration signaling itself, specifically JNK signaling, could induce the upregulation of *mol*. Canonical JNK signaling acts through the transcription factor AP-1, which is a heterodimer of Jun (FBgn0001291) and Fos (FBgn0001297)(121). Downstream genes are regulated through AP-1 binding to the conserved TPA-responsive element (TRE) sequence (TGAC/GTCA) (122). Indeed, there are three consensus TRE sites at the *mol* locus: one 2 Kb upstream of the transcription start site, one in the first intron, and one in the fifth intron (Fig 8A).

**Fig 8.**
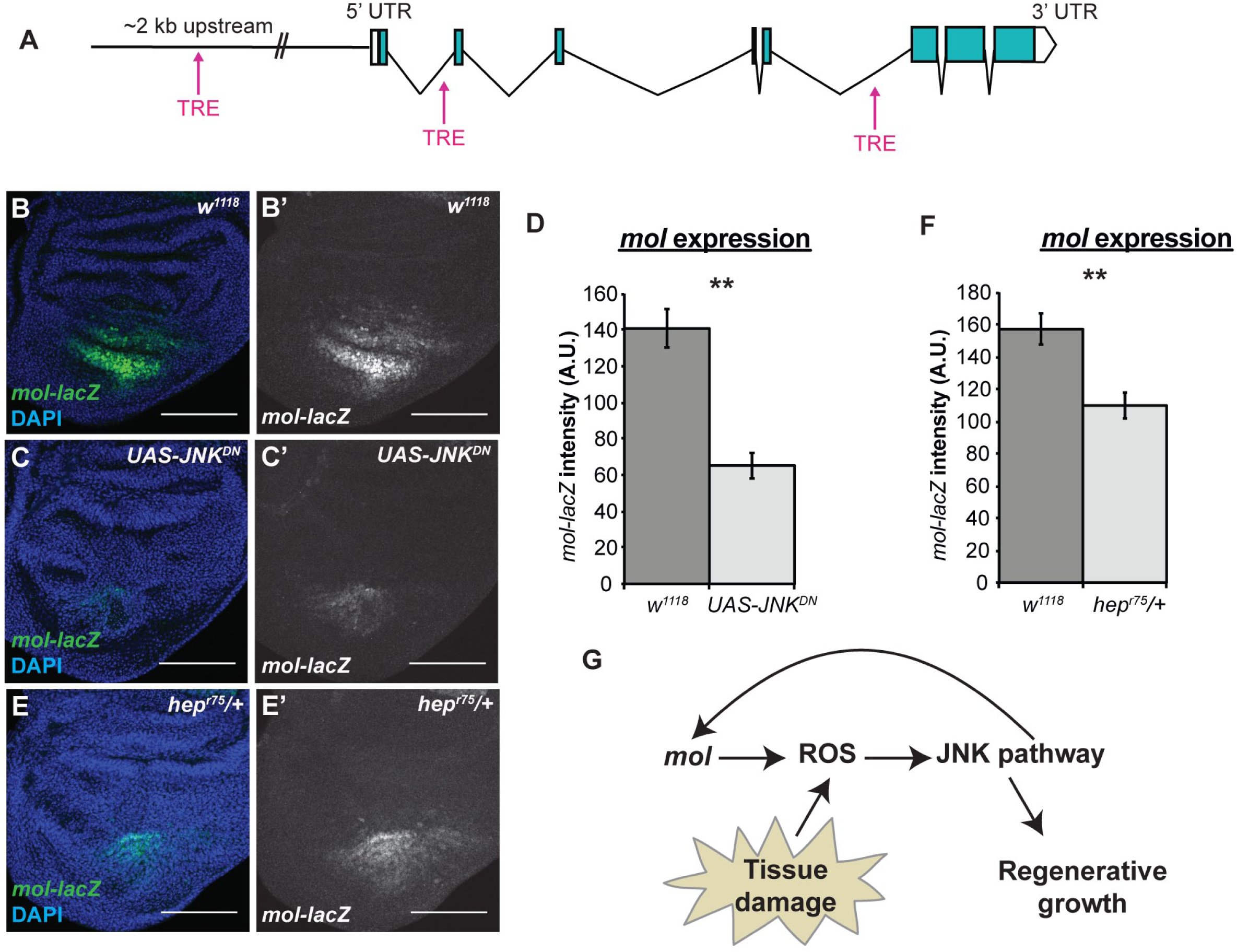
Expression of *mol* is regulated by JNK signaling. (A) Schematic of the *mol* locus showing the relative positions of three canonical TRE sites. (B,C) Anti-β-galacosidase immunostaining showing expression of the *mol-lacZ* (green) reporter in control (B) and *UAS-JNK^DN^* (C) R24 discs. (D) Quantification of *mol-lacZ* fluorescence from the immunostaining. *w^1118^* n=10 discs, *UAS-JNK^DN^* n=10 discs. (E) Anti-β-galacosidase immunostaining showing expression of the *mol-lacZ* reporter in a *hep^r75^*/+ R24 regenerating disc. (F) Quantification of *mol-lacZ* fluorescence from the immunostaining. *w^1118^* n=7 discs, *hep^r75^*/+ n=8 discs. Scale bars are 100 μm. Error bars are SEM. **p<0.002

To determine the extent to which JNK signaling is required for *mol* expression after tissue damage, we inhibited JNK signaling by expressing a dominant-negative JNK (*UAS-JNK^DN^*) (FBgn0000229)(123) under the control of *rn*-GAL4 during wing pouch ablation. Interestingly, expression of the reporter *mol-lacZ* was significantly decreased upon reducing JNK signaling through *UAS-JNK^DN^* Fig 8B-D), suggesting that JNK signaling is important for *mol* upregulation after tissue damage. To confirm this finding, we also examined *mol-lacZ* expression in regenerating wing discs that were heterozygous mutant for *hemipterous (hep)* (FBgn0010303), which encodes a JNK kinase (124). Expression of the *mol-lacZ* reporter was also significantly decreased in female *hep^r75^*/+ regenerating wing discs (Fig 8E,F). Thus, JNK regulation of *mol* expression constitutes a positive feedback loop that sustains JNK signaling.

ROS activates both JNK and p38a (FBgn0015765) in the regenerating wing disc (43). To determine whether p38a signaling also induces a positive feedback loop through *mol*, we examined *mol-lacZ* expression in regenerating discs that were heterozygous mutant for *p38a^1^*. Interestingly, reduction of p38a did not affect *mol-lacZ* expression (Fig S8). Thus, JNK signaling is required for upregulation of *mol* expression after tissue damage, which is in turn required for sustaining ROS production in the regenerating epithelium and maintaining JNK signaling during late regeneration (Fig 8G).

## DISCUSSION

This work describes generation of a transcriptional profile of actively regenerating tissue, made possible by our genetically induced tissue ablation system (6) and our technical advances enabling isolation of sufficient numbers of blastema cells (41). Through analysis of the expression data, followed by functional validation of differentially expressed genes, we have discovered a key positive feedback loop that uses JNK-induced upregulation of the Duox-maturation factor encoded by *mol* to sustain ROS production, JNK signaling, and late regeneration. Moreover, elevated ROS levels appear to be sustained in other regeneration models such as amputated zebrafish fins and *Xenopus* tails, where they promote signaling and the later stages of regenerative growth (99,100,102). Therefore, the positive feedback loop we have identified may facilitate long-term regeneration signaling in many animals.

This is the first report of upregulation of a Duox maturation factor as a key aspect of the regeneration response. Other cellular functions that are regulated by DUOXA/NIP have only recently been identified. For example, DUOXA/NIP affects differentiation in murine skeletal muscle myoblasts (125), murine thyroid hormone production and cerebellar development (126), and the response to bacterial infections in the murine gut (127), as well as development of the exoskeleton in *C. elegans* (128), and recruitment of hemocytes to wounds in the *Drosophila* embryo epidermis and neutrophils to airways in mice (129,130). Here we describe a role for *mol* during wing disc regeneration and show that while *mol* is transcriptionally upregulated, Duox levels do not change according to our transcriptional profile, indicating that fine-tuning of ROS levels can be achieved by changes in expression of the maturation factor rather than the enzyme itself. This regulative strategy may be deployed in many other cases in which ROS act as crucial signaling molecules.

In addition to the transcriptional changes observed in regulators of ROS, many of the other changes in gene expression can be combined with our current understanding of tissue regeneration to identify novel and interesting relationships between developmental genes and signals and tissue regeneration. For example, our data indicated downregulation of the hormone receptor Hr78 in regenerating tissue. The expression of Hr78 in the wing disc appeared to be in some of the pro-vein regions (Fig 3B). Tissue damage in the wing disc leads to a transient loss of cell-fate gene expression, including in the pro-veins, during regeneration (6,94). Thus, Hr78 may be a novel wing vein fate gene whose expression is downregulated along with the other known vein fate genes after tissue damage.

As an additional example, we observed differential regulation of various nuclear hormone receptor genes that are transcriptionally regulated by the hormone ecdysone (131). Regenerating animals delay metamorphosis to accommodate regrowth of the damaged tissue by regulating ecdysone signaling, which controls developmental transitions (132). Ecdysone targets that we found downregulated in regenerating wing discs include *Hormone receptor 46* (*Hr46/Hr3*) (FBgn0000448), *Hormone receptor 4* (*Hr4/CG42527*) (FBgn0264562), and *Ecdysone-induced protein 78C* (*Eip78C*) (FBgn0004865) (Table S2). Interestingly, we also see upregulation of *Cyp18a1* (FBgn0010383), a cytochrome P450 enzyme that exerts negative feedback regulation on ecdysone signaling by decreasing intracellular levels of ecdysone (133). Thus, *Cyp18a1* may be upregulated to ensure that ecdysone signaling stays low in the regenerating tissue to reinforce the developmental checkpoint induced by tissue damage.

Regeneration involves orchestration of various cellular processes to repair and replace the damaged body part. It requires coordination of proliferation, growth, patterning, and changes in cell architecture and movement in a highly regulated manner. These dramatic changes could be coordinated by key transcription factors. Several transcription factors are differentially expressed in our profile, including *chinmo*, *Ets21C*, *AP-2/TfAP-2* (FBgn0261953), *fru*, *Atf3*/*A3*-3, *dve* and *Blimp-1* (FBgn0035625). These transcription factors could lie at the center of regulatory networks that bring about key cellular changes. For example, Ets21C is a known downstream target of JNK signaling in wound healing (86), and EGFR signaling in the intestinal stem cells (134), and is also required as a co-factor for the JNK pathway transcription factor AP-1 in regulating transcriptional targets during tumor formation (84,85). Thus, its expression in the regenerating wing disc could result from integration of multiple signals, and its requirement in regeneration may be due to its role in promoting expression of JNK targets. Further investigation into the mechanisms of these transcription factors will lead to a better understanding of regeneration.

Regeneration is a tightly controlled process, requiring a balance between positive and negative regulators so that growth is stimulated but not deregulated. Indeed, our functional analysis demonstrated that several of the upregulated genes, including *heartless* and *Nox*, serve to restrict regeneration, as regeneration improved in heterozygous mutant animals. Therefore, functional analysis is critical for interpretation of gene expression data, as drawing conclusions based on differential expression alone can be misleading. Indeed, it was through functional analysis that we identified *mol*, and not *Nox*, as the critical regulator that promotes sustained ROS production and JNK signaling, completing the positive feedback loop that sustains regeneration. Further functional analysis of differentially expressed genes will likely reveal additional mechanisms that control tissue regeneration.

## Materials and Methods

### Ablation system

Tissue ablation was carried out as described previously (6,82) using *rnGal4*, *UAS-rpr*, and *tubGAL80^ts^* to regulate cell death spatially and temporally, with a thermal shift from 18° to 30° C for 24 hours during the early third larval instar. To synchronize development, eggs were collected for four hours on grape juice plates, first-instar larvae were collected shortly after hatching at two days after egg laying and transferred to vials, and the vials underwent the thermal shift at 7 days after egg laying, which was determined to be just after molting by counting mouth hooks.

### Fly lines

Flies were reared on standard molasses medium at 25° except during regeneration experiments. Following *Drosophila* lines were obtained from the Bloomington Stock Center or were gifts as noted: *w^1118^; rnGAL4, UAS-rpr, tubGAL80*^ts^/ *TM6B, tubGAL80* (Smith-Bolton et al., 2009), *w^1118^; rnGAL4, UAS-rpr, tub-GAL80*^ts^/TM6B; *y^1^,w^*^; Mi{MIC}nub^MI05126^* (BL37920)(48), *y^1^,w^67c23^; P{lacW}chinmo^k13009^/CyO* (BL10440)(135), *y^1^, w^*^ Mi{MIC}pigs^MI11007^* (BL56274)(48), *P{PZ}Alp4^07028^, ry^506^* (BL12285)(135), *w^1118;^ PBac{Ets21CGFP.FLAG}VK00033/TM3, Sb^1^* (BL38639), *P{PZ}osp^00865^; ry^506^ P{PZ}zfh1^00856^/TM3, ry^RK^ Sb^1^ Ser^1^* (BL11515)(136), *y^1^ w^67c23^; P{lacW}mol^k11524a^/CyO* (BL12173)(135), *ry^506^ P{PZ}fru^3^/MKRS* (BL684)(135), *w;;pBAC[atf3::EGFP]/TM6B* (gift from M. Uhlirova)(137), *nlaZ:GFP[R2]* (gift from M. Ganfornina)(69), *w^1118^; P{10xStat92E-GFP}1* (BL26197)(66), *cn^1^ P{PZ}dve^01738^/CyO; ry^506^* (BL11073)(135), *cn^1^ P{PZ}sm^05338^/CyO; ry^506^* (BL11403)(78), *y^1^ w^*^; P{PTT-GB}LamC^CB04957^ ttv^CB04957^/SM6a* (BL51528)(138), *y^1^ w^*^; Mi{PT-GFSTF.1}AdoR^MI01202-GFSTF.1^/TM6C, Sb^1^ Tb^1^* (BL60165)(139), *y^1^ w^*^; P{lacW}Thor^k13517^* (BL9558)(67), *y^1^ w^*^; Mi{PT-GFSTF.0}kay^MI05333-GFSTF.0^* (BL63175)(139), *w^1118^; PBac{corto-GFP.FPTB}VK00037* (BL42268), *w^1118^; PBac{Hr78-GFP.FLAG}VK00037* (BL38653), *NC2β-GFP* (BL56157), *w^1118^;PBac{NC2β-GFP.FPTB}VK00033* (BL60212)(139), *Hml*Δ*RFP* (gift from K. Bruckner)(115), *TRE-red* and *gstD-GFP* (gifts from D. Bohmann)(114), *Ets21C^f0369^* (BL18678)(140), *y^1^ w^*^; Mi{MIC}CG9336^MI03849^* (BL36397)(139), *y^1^ w^67c23^; P{lacW}Col4a1^K00405^/CyO* (BL10479)(135), *y^1^ w^67c23^; P{lacW}vkq^k00236^* (BL10473)(135), *P{PZ}Thor^06270^ cn^1^/CyO; ry^506^* (BL11481)(141), *w^1^; P{UASSod1.A}B36* (BL24754), *w^1^; P{UAS-CatA}2* (BL24621), *w^1^; P{UASSod2.M}UM83* (BL24494), *w^1118^; PBac{RB}mol^e02670^/CyO* (BL18073)(140), *y^1^ sc^*^ v^1^; P{TRiP.HMS02560}attP40 (UAS-mol^RNAi^)* (BL42867), *y^1^ v^1^; P{TRiP.GL00678}attP40* (*UAS*-*Duox^RNAi^*) (BL38907), *y^1^ v^1^; P{TRIP.GL00678}attP40 (UAS*-*Nox^RNAi^)* (BL32902), *y^1^ w^*^; Mi{MIC}Nox^MI15634^/SM6a* (BL61114)(139), *puc^E69^* (Martin-Blanco et al., 1998), *UAS*-*JNK^DN^* (142), *w*^*^ *hep^r75^/FM7C* (BL6761)(124), *w^*^; P{neoFRT}82B p38a^1^* (BL8822)(143).

### Immunohistochemistry and microscopy

Immunostaining was carried out as previously described (6). Anitbodies and dilutions used were Anti-Nubbin (1:500) (gift of S. Cohen) (47), mouse anti-βgal (1:100) (DSHB; 40-1a-s), rabbit anti-βgal (1:500) (MP Biomedicals), mouse anti-GFP (1:10)(DSHB 12E6), rabbit anti-Myc (1:500) (Santa Cruz Biotech d1-717 sc-28207), rabbit anti-PH3 (1:500) (Millipore), mouse anti-Nimrod (1:1000)(gift from I. Ando)(116) and anti-Twist (1:200) (gift from A. Stathopoulos)(144).

Alexa Fluor (AF) secondary antibodies from Molecular Probes were AF488, AF555 and AF633 (used at 1:500). Nuclei were labeled with DAPI (Sigma)(1:5000).

EdU incorporation was carried out using the click-it EdU Alexa Fluor 594 Imaging kit (Molecular Probes) as previously described (145). Samples were mounted in Vectashield (Vector Labs).

Immunostained samples were imaged on a Zeiss LSM 700 confocal microscope and images were processed using ZenLite, Adobe Photoshop and Image J software. Bright-field imaging of adult wings was done on an Olympus SZX10 microscope using the CellSens Dimension software, and images were processed using Image J.

### ROS detection

ROS were detected in imaginal discs using Dihydroethidium (DHE) (D11347, Molecular Probes) using the protocol described in Owusu-Ansah et al. (146), with slight modifications. Briefly, larvae were dissected in Schneider’s medium (SM). DHE was reconstituted in DMSO and then added to SM at a concentration of 30nM. Samples were incubated in this DHE solution for 5 minutes (mins) on a shaker followed by three quick washes in SM. The samples were then fixed in 7% paraformaldehyde made in 1X phosphate buffer saline (PBS) for 7 mins. Samples were rinsed once in 1X PBS and imaginal discs immediately dissected out to mount in Vectashield with DAPI. The samples were imaged on the confocal immediately to avoid oxidation of the DHE by the environment.

### Data quantification and statistical analysis

Fluorescence intensity analysis was performed using single confocal slices. Average intensity was calculated by measuring intensity values in three equal-sized boxes in the pouch region of the wing disc in Image J, except for the *gstD-GFP,* whose expression was not uniform and thus was quantified by measuring GFP intensity in the entire pouch area. Average intensities of multiple wing discs were combined to calculate the final average intensity plotted in the graphs. For measuring the pouch area, a maximum projection of all the confocal slices was taken and the Nubbin-expressing area measured in Image J. Graphs were plotted in Excel, R and GraphPad Prism 7.0.

For imaginal disc measurements and immunofluorescence quantifications, the Welch’s t-test was performed using R and GraphPad Prism 7.0. For the adult wing size assay, chi-squared tests were performed using GraphPad online tools. Statistical analyses for adult wing measurements were performed using Welch’s t-test.

### Blastema cell isolation and RNA library preparation

The ablated regenerating discs had the genotype *nub-GFP/+; rn-Gal4, GAL80^ts^, UAS-rpr/+,* while the mock-ablated controls had the genotype *nub-GFP/+; rn-Gal4, GAL80^ts^*/+. Cells were isolated for the transcriptional profile as previously described (41). Briefly, discs were dissected using teams of 4 researchers dissecting simultaneously to maximize the number of discs obtained per sample. TrypLE Select (Life Technologies) was used to achieve rapid dissociation of the disc cells. The GFP+ cells were sorted via FACS. mRNA from the isolated cells was prepared using an RNeasy Mini Kit (#74104, Qiagen). Multiple days of dissections and RNA preparation were pooled such that each biological replicate consisted of approximately 600 regenerating imaginal discs, 86,000 GFP+ cells, and up to 900ng RNA. The undamaged controls consisted of 120 discs per replicate, which produced approximately 106,500 GFP+ cells and 1000ng RNA. The accuracy of the sorting was previously confirmed (41). RNA quality was confirmed using a Bioanalyzer (Agilent 2100). Library generation was carried out using Illumina’s TruSeq Stranded RNA Sample Prep kit. Sequencing was carried out on a HiSeq2000 using a TruSeq SBS sequencing kit version 3. The Roy J. Carver Biotechnology Center at the University of IIlinois at Urbana-Champaign performed the library preparation and sequencing.

### Bioinformatics

Fastq reads were trimmed using FASTQ Quality Trimmer (v.1.0.0) and adaptor sequences were removed using Clip (v.1.0.1) in Galaxy (147). Paired-end reads were aligned through Tophat2 (v.0.6) (51,52) against the *Drosophila melanogaster* genome (NCBI, build 5.41) with a maximum of 2 mismatches permitted. Intron length was set between 20 and 150,000, and a gene model was provided as GTF (NCBI, build 5.41). FPKM estimation was done using Cufflinks (v.0.0.7) (51), and both bias-correction and multi-read correction were performed. Differential expression analysis was performed using Cuffdiff (51), geometric library normalization was performed and the False Discovery Rate was set at 0.5. Furthermore, aligned reads were counted using HTSeq (v.0.3.2)(51). All bioinformatics analysis was performed using the Galaxy suite (147).

## Acknowledgments

The authors would like to thank A. Brock, K. Schuster, and L. O’Brien for critical reading of the manuscript and helpful discussions, D. Bohmann, K. Bruckner, A. Stathopoulos, I. Ando, M. Uhlirova, M. Ganfornina, and S. Cohen, the Bloomington *Drosophila* Stock Center (NIH P40OD018537), the TRiP project at Harvard Medical School (NIH/NIGMS R01-GM084947), the Vienna *Drosophila* Resource Center, and the Developmental Studies Hybridoma Bank (NICHD, University of Iowa) for reagents. The authors would also like to thank B. Pilas at the Flow Cytometry Facility, A. Hernandez at the High-Throughput Sequencing and Genotyping Unit, and R. Khetani at the High Performance Biological Computing Center, all of which are part of the Roy J. Carver Biotechnology Center at the University of Illinois Urbana-Champaign.

## Author contributions

The experiments were conceived and designed by S.J.K. and R.K.S.-B. The experiments were performed by S.J.K., S.N.F.A., A.S., and Y.T. The manuscript was prepared by S.J.K. and R.K.S.-B.

